# A small-molecule fluorescent probe for live-cell imaging of endocytosis

**DOI:** 10.1101/2021.07.29.454398

**Authors:** Ryo Seino, Hidefumi Iwashita, Masataka Takahashi, Takatoshi Ezoe, Munetaka Ishiyama, Yuichiro Ueno

**Affiliations:** Dojindo Laboratories, Kumamoto, Japan

**Keywords:** Endocytosis, endosome, lysosome, live-cell imaging

## Abstract

Endocytosis involves plasma membrane-derived vesicles for the recycling of intra- and extracellular components. Increasing evidence suggests that endocytosis is related to maintaining intracellular homeostasis and defense against disease. Consequently, investigation of the endocytic pathway attracts considerable scientific interest. This study reports live-cell imaging of endocytosis using the newly-developed fluorescent probe ECGreen. We demonstrate that ECGreen is not membrane permeable and its fluorescence signal increases in acidic conditions. Because of these characteristics, ECGreen remains on the plasma membrane, and then shows increased fluorescence when it is internalized into the acidic vesicles formed in the endocytic process. ECGreen allows direct observation of the internalized vesicle; it is a valuable new probe for endocytic imaging.

## Introduction

Endocytosis involves macromolecules being taken up into plasma membrane-derived vesicles [1]. The vesicles involved in the endocytotic pathway, including endosomes (early endosomes and late endosomes) and lysosomes, are acidic and are reported to interact with other organelles [2-3]. Through these interaction, the endocytic pathway contributes to maintaining intracellular homeostasis [4-5]. Moreover, recent findings reveal that disruption of endocytosis is closely related to certain neurodegenerative disorders [6-7]. Accordingly, investigation of the endocytic pathway has attracted a great interest in the scientific fields.

Visualizing endocytic vesicles helps investigation of the endocytic pathway, and understanding of the interactions of the endocytic vesicles with other organelles and their roles in cellular functions. Development of a reliable and convenient method for monitoring the intracellular dynamics of the endocytic pathway is necessary. The most famous method for detecting endocytic vesicles is immunofluorescence staining of specific maker proteins. Immunofluorescence staining of early- or late-endosomal membrane marker proteins, Rab5 and Rab7 respectively, is often used in co-localization studies to confirm whether proteins of interest are associated with the endocytic pathway [8-9]. However, this method cannot be applied in live-cell imaging of endocytic vesicles because cell fixation and permeabilization are necessary. Rab5 and Rab7 tagged with fluorescent protein have been used to detect the dynamics of endocytic vesicles in live cells. The application of this approach is, however, limited to certain types of cells because of the requirement for plasmid transfection for protein expression [10-11,13]. Fluorescent dextran conjugates, internalized via endocytosis, are widely used for monitoring the endocytic pathway in living cells [12-13]. Fluorescent dextran conjugates are small-molecule fluorescent probes, and can be applied in any type of cell. However, the size of the dextran conjugate affects both internalization and dynamics in the endocytic pathway [14], suggesting that observations made using dextran conjugates should be treated with caution. Therefore, a new, reliable tool for monitoring the intracellular dynamics of endocytosis is needed. Here, we report live-cell imaging of endocytosis using the novel probe ECGreen.

## Materials and Methods

### 1. Reagents and Instruments

All reagents and buffers were purchased from Fujifilm Wako Pure Chemical Corporation, unless otherwise noted. ECGreen was obtained from Dojindo Laboratories. LysoTracker Red DND-99 (Thermo Fisher Scientific) was used for staining lysosomes. Cell Light-Red Fluorescent Protein (RFP) reagents (Thermo Fisher Scientific) were used for staining the early endosome, late endosome, and lysosome. pHrodo Green Dextran, 10,000 MW for Endocytosis (Thermo Fisher Scientific) was used for visualizing the endocytic pathway. All fluorescent dyes were used following the product manuals. Fluorescence spectra were measured on an FP-6300 fluorescence spectrophotometer (JASCO). Fluorescence images were obtained by LSM 800 confocal laser scanning microscopy (Zeiss), with excitation at 405 nm (for ECGreen), 488 nm (for pHrodo Green Dextran), and 561 nm (for LysoTracker Red and RFP), using a 500–550-nm filter for ECGreen and pHrodo Green Dextran, or a 550–650-nm filter for LysoTracker Red and RFP.

### 2. Experimental Procedure

#### Spectrophotometric Measurement

The pH-dependent fluorescence intensity of ECGreen was measured in 50% dimethylsulfoxide in MES at pH 4.0, 4.5, 5.0, 5.5, 6.0, 6.5, 7.0, 7.5, 8.0 and 8.5, using a fluorescence spectrophotometer (excitation 405 nm, emission 520 nm).

#### Cell Culture

HeLa cells were cultured in Minimum Essential Medium (Thermo Fisher Scientific) supplemented with 10% (v/v) fetal bovine serum, 1% L-glutamine, 1% non-essential amino acids, 1% penicillin, and 1% streptomycin. Cells were maintained in a humidified 5% CO_2_ incubator at 37 °C.

## Results and Discussion

ECGreen, a non-membrane-permeable small-molecule fluorescent dye, is internalized via endocytosis and its fluorescence intensity *in vitro* is enhanced by increasing acidity (Figure 1A and 1B). Having these characteristics, ECGreen may allow direct observation of the dynamics of endocytic vesicles in live cells.

**Figure 1.**
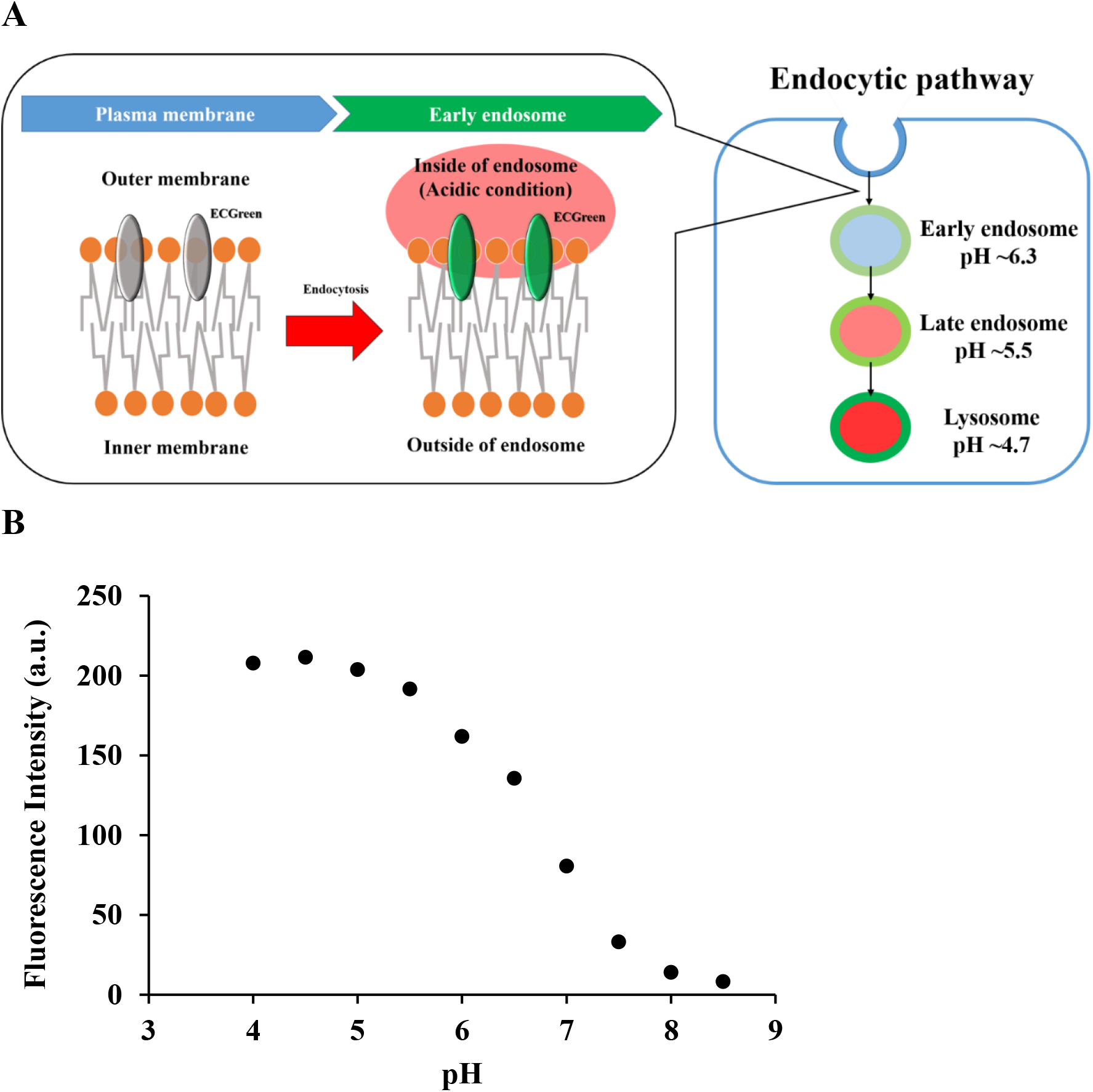
Characteristics of ECGreen. (A) Scheme illustrating the principle of detection of endocytosis using ECGreen. (B) pH-dependent changes of the fluorescence intensity of ECGreen (1.0 μM) in buffer solutions (pH 4.0 – 8.5), excited at 405 nm (emission 520 nm). A working solution of ECGreen (10 mM in DMSO) was diluted with MES buffer.

To address whether ECGreen was internalized via endocytosis, the uptake of ECGreen was firstly investigated with low temperature incubation. Since endocytosis is a temperature dependent process, cells maintained at low-temperature are often used as a reference for the absence of endocytosis [15]. As shown in Figure 2A, fluorescent puncta of ECGreen were observed and co-localized with lysosomes stained by LysoTracker Red in HeLa cells incubated at 37 °C, whereas the uptake of ECGreen was dramatically inhibited at 4 °C. In contrast, fluorescent puncta of LysoTracker Red, a membrane-permeable dye, was observed under low temperature condition. These results revealed that ECGreen is internalized not directly through the cellular membrane but via endocytosis.

**Figure 2.**
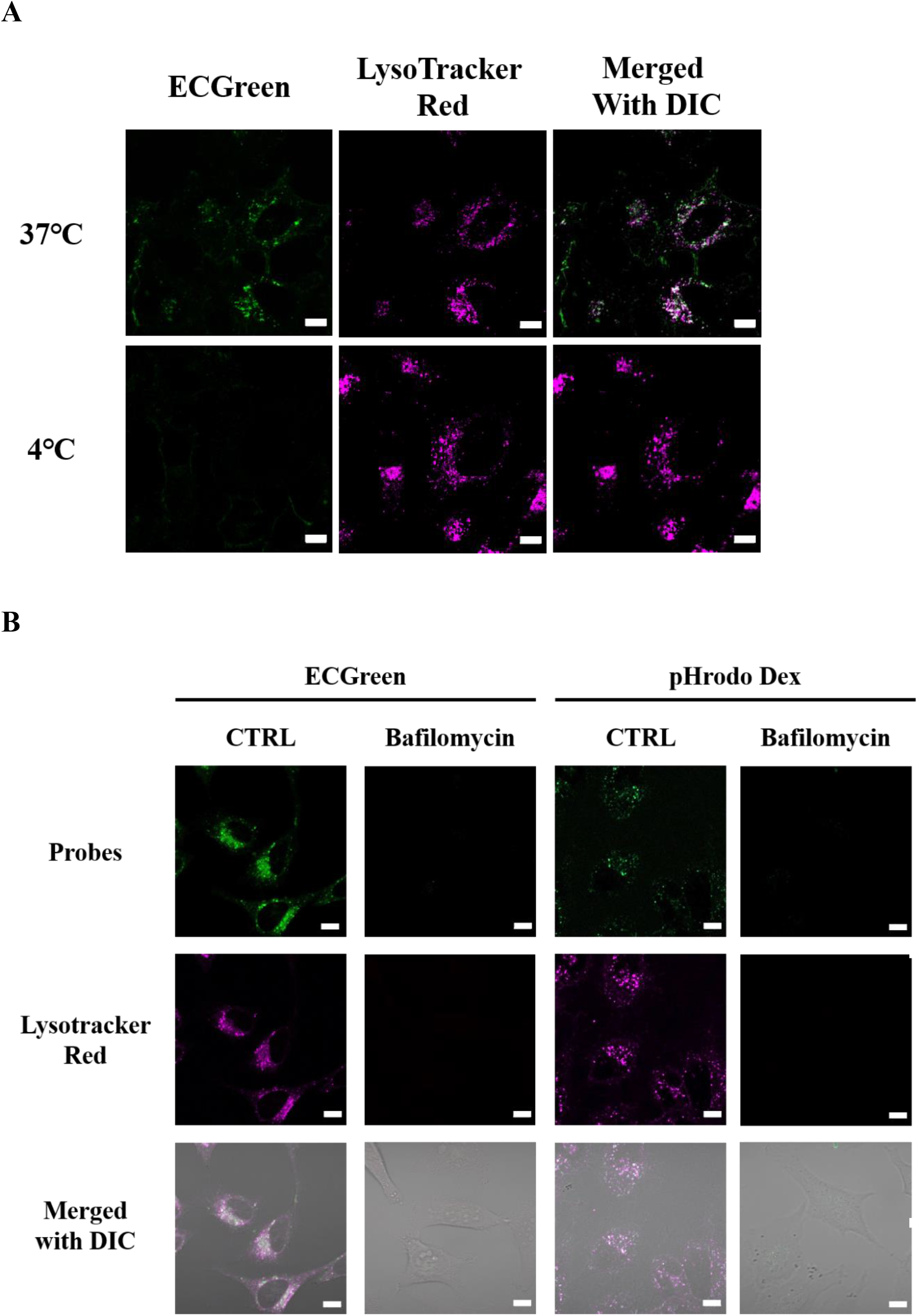
Internalization of ECGreen via endocytosis. (A) HeLa cells were stained with 5 µM ECGreen and 100 nM LysoTracker Red in Minimum Essential Medium containing 10% fetal bovine serum and incubated for 30 min at 37 or 4 °C. After washing with Hanks’ Balanced Salt Solution (HBSS), the cells were observed under a fluorescence microscope. (B) HeLa cells were pretreated with 100 nM bafilomycin for 30 min before staining with 5 µM ECGreen or 50 µg/ml pHrodo Green Dextran (pHrodo Dex) in the presence of 100 nM LysoTracker Red. After 30 min of incubation, the cells were washed and observed under a fluorescence microscope. Bars: 10 µm.

We also verified the effect of endo-lysosomal acidification on the fluorescence of ECGreen. Endocytic organelles and lysosomes are acidic, and this is important for endosome maturation [16]. Dextran conjugated to the pH-sensitive dye pHrodo Green is a well-known tool for monitoring acidification of the endocytic pathway. Like ECGreen, the fluorescence intensity of this dye increases in response to acidic pH [12-13]. Staining results (Figure 2B) showed that the fluorescent puncta of pHrodo Green Dextran were co-localized with lysosomes. In contrast, no fluorescent puncta of either ECGreen or pHrodo Green Dextran were observed when cells were treated with bafilomycin A1, an inhibitor of endo-lysosomal acidification. Our data suggest that ECGreen is internalized via endocytosis and its fluorescence intensity is enhanced when it is internalized into the acidic vesicles formed in the endocytic process.

Next, we verified the intracellular localization of ECGreen. Co-localization studies (Figure 3) revealed that ECGreen co-localized with the early endosomal marker Rab5, the late endosomal marker Rab7, and the lysosomal marker LAMP1, each tagged with RFP. Therefore, ECGreen co-localizes with endocytic organelles and lysosomes.

**Figure 3.**
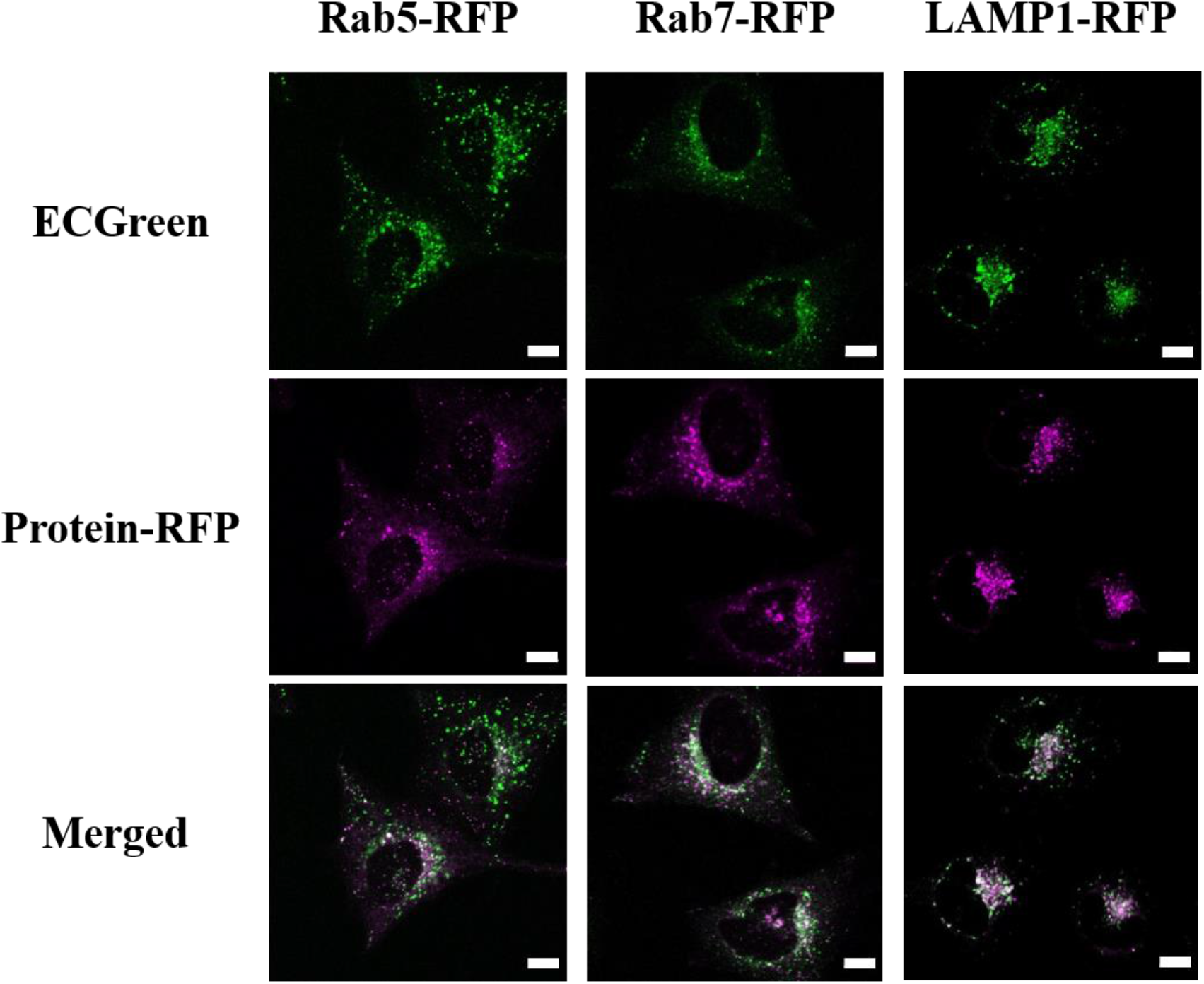
Intracellular localization of ECGreen. HeLa cells transiently expressed Rab5–RFP (a marker for early endosomes), Rab7–RFP (for late endosomes), or LAMP1–RFP (for lysosomes) using Cell Light Reagent-Red Fluorescent Protein (RFP). RFP-expressing HeLa cells were stained with 5 µM ECGreen for 30 min. After washing with HBSS, the cells were observed under a fluorescence microscope. Bars: 10 µm.

To determine whether ECGreen localization depends on endosomal membrane trafficking, its localization was examined in conditions where endosomal maturation was inhibited. Phosphatidylinositol 3-phosphate (PI3P), produced from phosphatidylinositol by class III PI3-kinase, is predominantly localized to early endosomes, and controls endosome maturation [17-18]. Consequently, wortmannin, a potent inhibitor of class I and III PI3-kinases, blocks maturation from early to late endosomes [19-21]. In the presence of wortmannin, RFP–Rab5 showed a ring-shaped structure, indicating a swollen endosome phenotype and location at the endosomal membrane (Figure 4A, upper panel, white arrow). In cells treated with wortmannin, similar observations were made for ECGreen (Figure 4A, lower panel, white arrow); its ring structure co-localized with that of RFP–Rab5 (Figure 4B, upper panel, white arrows). However, RFP–Rab7 and RFP– LAMP1 did not co-localize with the wortmannin-induced ring structure of ECGreen (Figure 5A and 5B). On the other hand, pHrodo Green Dextran showed large puncta, but not a ring shape, and was located inside of the Rab5-ring structure in the presence of wortmannin (Figure 4B, lower panel, white arrowheads), indicating that fluorescent dextran conjugates stain internal space of endocytic vesicle. These findings indicate that ECGreen specifically locates to the early endosomal membrane and is then transported to the later endocytic pathway.

**Figure 4.**
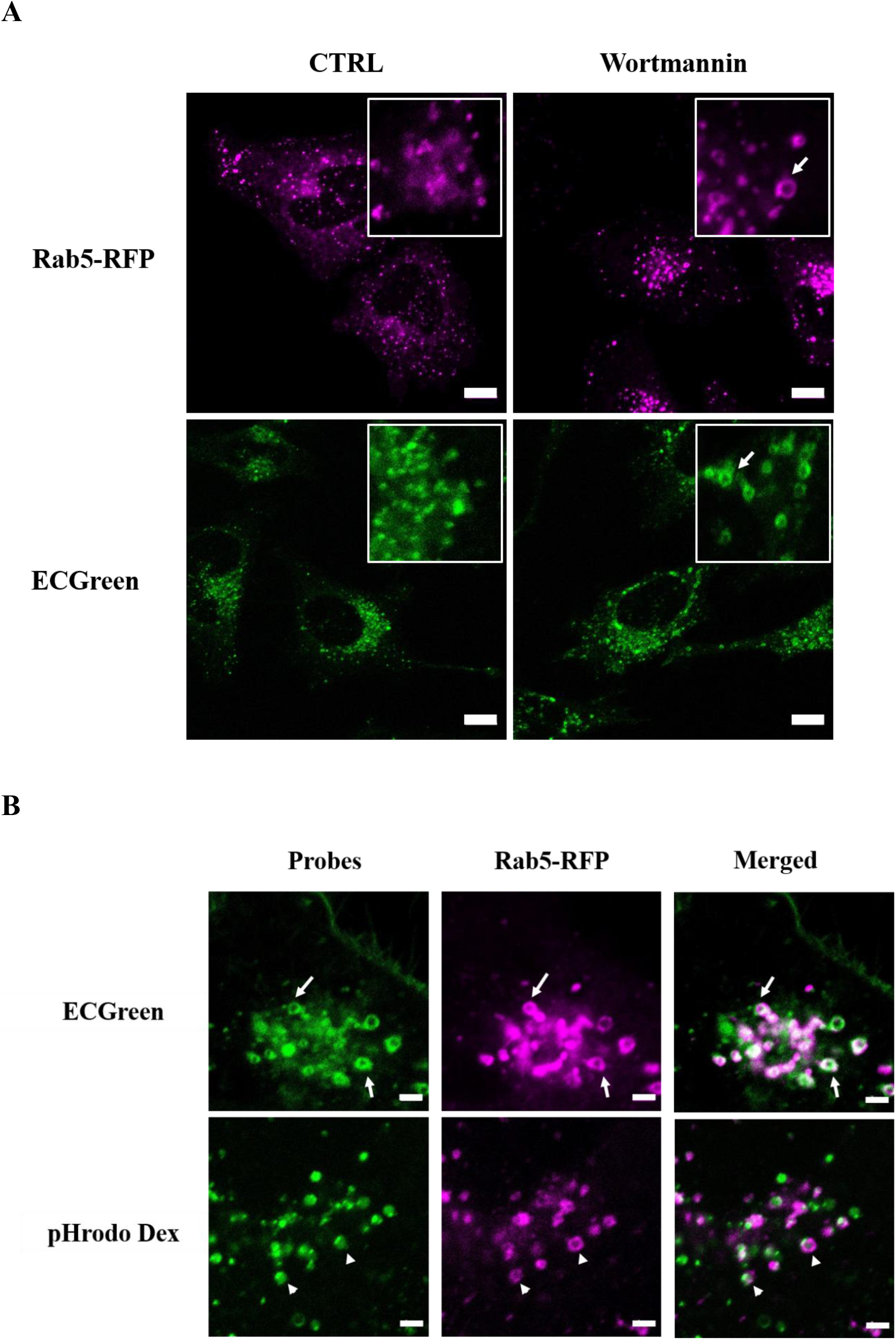
Localization of ECGreen to the early endosomal membrane. (A) Rab5–RFP expressing HeLa cells were treated with 100 nM wortmannin for 30 min with or without 5 µM ECGreen. After washing with HBSS, the cells were observed under a fluorescence microscope. Arrows: membrane-like ring-structure signals. Bars: 10 µm. (B) Rab5–RFP-expressing HeLa cells were pretreated with 100 nM wortmannin for 30 min before staining with 5 µM ECGreen or 50 µg/ml pHrodo Green Dextran for 30 min. After washing with HBSS, the cells were observed under a fluorescence microscope. Arrows and arrowheads respectively indicate overlapping ECGreen/Rab5–RFP ring-structure signals and pHrodo Dex surrounded by Rab5–RFP. Bars: 2 µm.

**Figure 5.**
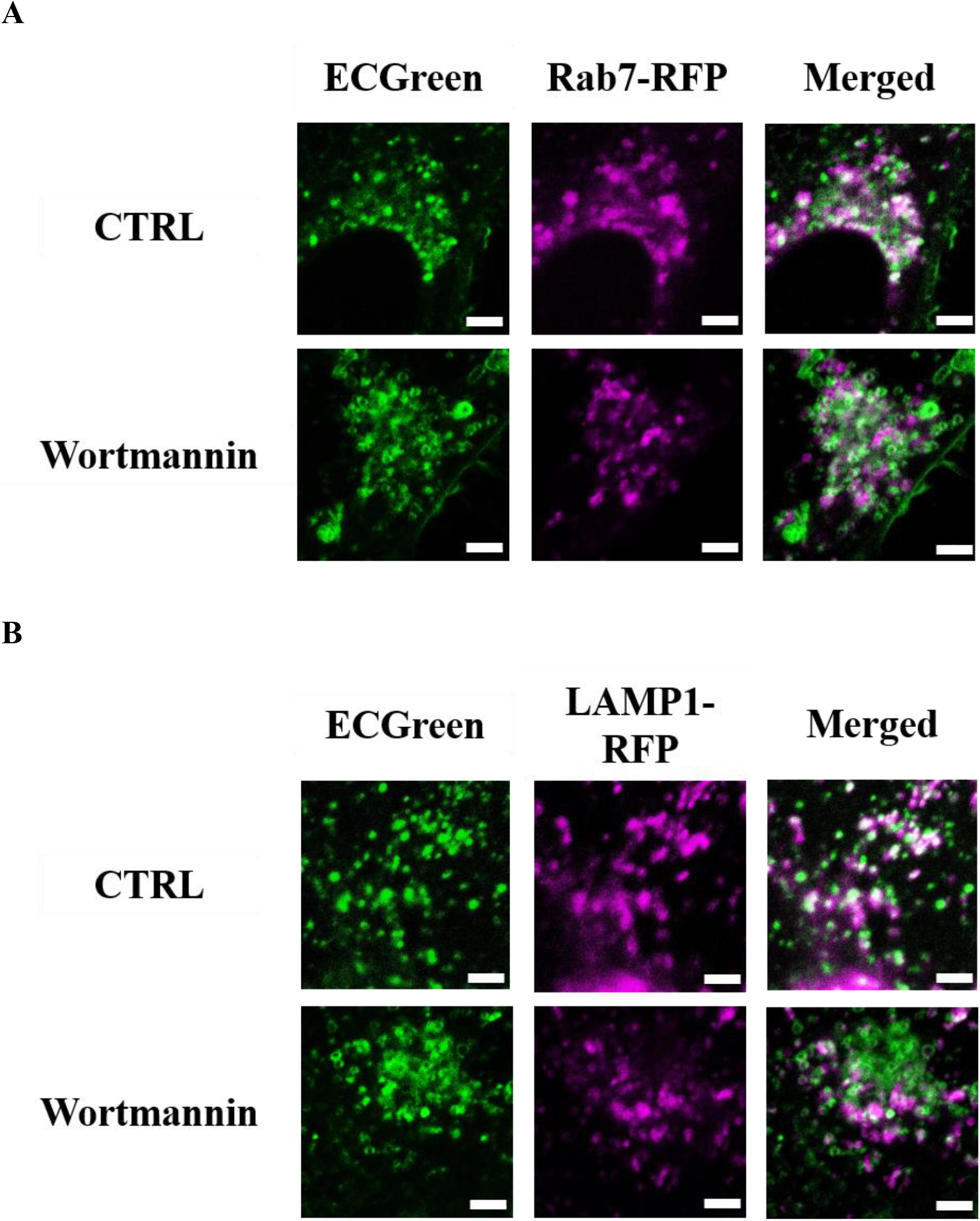
Transport of ECGreen along the endocytic pathway. HeLa cells expressing Rab7–RFP (A) or LAMP1–RFP (B) were pretreated with 100 nM wortmannin for 30 min. Then, 5 µM ECGreen was added to the cells. After 30 min of incubation, the cells were washed and observed under a fluorescence microscope. Bars: 2 µm.

## Conclusion

Much remains to be elucidated in the field of endocytosis, and there is a need for reliable and convenient methods for live-cell monitoring of the endocytic pathway. ECGreen is a small-molecule fluorescent probe that is internalized via endocytosis and its fluorescence intensity is enhanced as acidity increases. Unlike conventional dextran conjugates, ECGreen allows direct observation of the membrane of an internalized vesicle in live cells. In future, this probe will be valuable for studying the molecular mechanisms and dynamics of endocytosis.

## Abbreviation

RFP: red fluorescent protein.

## Acknowledgement

We thank James Allen, DPhil, from Edanz Group (https://en-author-services.edanzgroup.com/ac) for editing a draft of this manuscript.

